# A 2D convolutional neural network for taxonomic classification applied to viruses in the phylum *Cressdnaviricota*

**DOI:** 10.1101/2023.05.01.538983

**Authors:** Ruither A. L. Gomes, F. Murilo Zerbini

## Abstract

Taxonomy, defined as the classification of different objects/organisms into defined stable hierarchical categories (taxa), is fundamental for proper scientific communication. In virology, taxonomic assignments based on sequence alone are now possible and their use may contribute to a more precise and comprehensive framework. The current major challenge is to develop tools for the automated classification of the millions of putative new viruses discovered in metagenomic studies. Among the many tools that have been proposed, those applying machine learning (ML), mainly in the deep learning branch, stand out with highly accurate results. One ML tool recently released that uses k-mers, VirusTaxo, was the first one to be applied with success, 93% average accuracy, to all types of viruses. Nevertheless, there is a demand for new tools that are less computationally intensive. Viruses classified in the phylum *Cressdnaviricota*, with their small and compact genomes, are good subjects for testing these new tools. Here we tested the usage of 2D convolutional neural networks for the taxonomic classification of cressdnaviricots, also testing the effect of data imbalance and two augmentation techniques by benchmarking against VirusTaxo. We were able to get perfect classification during k-fold test evaluations for balanced taxas, and more than 98% accuracy in the final pipeline tested for imbalanced datasets. The mixture of augmentation on more imbalanced groups and no augmentation for more balanced ones achieved the best score in the final test. These results indicate that these architectures can classify DNA sequences with high precision.

## INTRODUCTION

Taxonomy, defined as the categorization of objects or beings in an organized, hierarchical way, is a core part of science. Taxonomy allows scientists to better communicate with each other by linking objects or beings to stably named categories (taxa) (Gorbalenya *et al*., 2019). Viruses are probably the most diverse and ubiquitous biological entities on the planet (Paez-Espino *et al*., 2016). Viral discovery during most of virology’s history was based on purifying viruses from infected hosts or cell cultures and on PCR-based sequencing of viral genomes, with taxonomic assignments being made individually by different research groups for each new virus based on phenotypic characteristics of the infection and molecular properties (Datta *et al*., 2015). As stated by Fauquet (1999), “As in other biological systems, virus classification is an approximate and imperfect exercise. Like any other type of classification, it is a totally artificial and human-driven activity”. In recent years, major changes in the taxonomic framework were made towards a more direct genome-based classification (Simmonds *et al*., 2023).

As a consequence of the exponential increase in viral sequence data from high-throughput sequencing (HTS) of metagenomic samples, the International Committee on Taxonomy of Viruses (ICTV) decided, in 2017, to start accepting metagenomic generated sequences (after checking for completeness and quality) as true representatives of viruses (Simmonds *et al*., 2017). Since then, several taxa (species, genera, families and one order) have been created based exclusively on metagenomic-derived sequences (Koonin & Yutin, 2020; Krupovic *et al*., 2016). Additionally, the ICTV invited the virology community to assist in the development of new automated classification methods to deal with large datasets (Simmonds *et al*., 2017).

Different computational classification tools have been proposed in the two last decades to help dealing with the new reality in viral biodiversity. Most of these tools are based on sequence clustering methodologies using demarcation thresholds that separate taxonomic levels using sequence divergence between one or more conserved genes in a group. These methods are usually applicable for classification at the family and genus ranks (Pappas *et al*., 2021).

Machine learning (ML) algorithms have already lead to incredible developments for different types of data analyses. In bioinformatics, they have been successfully applied to a broad range of fields, from the prediction of RNA or protein structure to phylogenetic and biological network analyses. In taxonomy, ML methods are mostly based on extraction and selection of different features from datasets of genomic sequences, and development of prediction models that can identify the taxonomic rank of new sequences without the need for direct comparison with all members of a selected group using sequence alignments, as done by most other tools. Thus, ML algorithms are much less computationally intensive than sequence alignment-based methods, allowing the trained ML models to be used in huge query datasets than would be unfeasible otherwise (Auslander *et al*., 2021).

When dealing with text, audio, image, and video recognition, convolutional neural networks (CNNs), a branch of deep learning (DL), are powerful tools due to their ability to extract patterns from large input datasets (Li *et al*., 2022). In recent studies proposing new bioinformatic tools, CNNs have been applied with highly accurate results in some fields, such as the identification of enhancer-promoter interactions (Min *et al*., 2021), the prediction of DNA-binding proteins (Barukab *et al*., 2022; Zhang *et al*., 2019), and the detection of patterns in DNA methylation (Zeng & Gifford, 2017). Specifically for viral classification, some examples are Viral Genome Deep Classifier, DeepVirFinder, EdeepVPP and EdeepVPP-hybrid (Dasari & Bhukya, 2022; Fabijańska & Grabowski, 2019; Ren *et al*., 2020), and BERTax, a method that uses natural language processing and achieved 95% accuracy at the phylum level of all four “superkingdoms” defined by the authors (archaea, bacteria, eukaryota, and viruses) (Mock *et al*., 2022). With the exception of BERTax, the algorithms achieved highly accurate training results only in specific datasets. Two of them (DeepVirFinder and EdeepVPP) were only able to predict if the input was a viral sequence, not inferring actual taxonomic classification, and the third one (Viral Genome Deep Classifier) was only trained to classify subtypes of five human viruses that had more than 10 similar sequences available. And even in the case of BERTax, in spite of its good accuracy at the phylum level, classification at the genus level achieved only 75% accuracy, which is inferior to other tools. Another common aspect of these tools is the use of text techniques, such as BioVec (Asgari & Mofrad, 2015) or BERT (Devlin *et al*., 2018), as done in BERTax, as the representation of the genomic sequence. Moreover, most of the times these were applied to one-dimensional (1D) CNNs, with only a few works using two-dimensional (2D) CNNs, and fewer applying to the taxonomic classification.

For classification into taxonomic ranks, ML algorithms have been developed that use assembled contigs as inputs to classify the sequence with high accuracy. The most recently developed, VirusTaxo (Raju *et al*., 2022), is based on a simple but memory-intensive detection of unique sequence fragments of length *k* (k-mers). VirusTaxo was the first tool trained with almost all virus genera available (of both RNA and DNA viruses), removing only those that had unique sequences (named singletons). The algorithm obtained an average accuracy of 93% during training. VirusTaxo was benchmarked against two commonly used tools based on k-mers, Kraken2 (Wood *et al*., 2019) and CLARK (Ounit *et al*., 2015), and also against DeepVirFinder, a CNN-based viral sequence predictor. It got similar results as DeepVirFinder for the identification of viral contigs in metagenome datasets, while Kraken2 and CLARK detected almost 100 times fewer viral sequences in the tested samples. Also, VirusTaxo achieved similar results as the other two k-mer-based tools while predicting taxonomic classification at the genus level from a dataset of partial and fully assembled contigs from the virus SARS-CoV-2.

The existing tools have different limitations for the identification and taxonomic classification of viral sequences. To date, only VirusTaxo can achieve the identification of viral sequences in metagenomic datasets and also predict the taxonomic classification of those sequences with confidence. However, this tool also has limitations, such as high RAM usage (for common computers, >24 GB) and the fact that k-mer-based approaches may be too sensitive to data-specific characteristics such as mutation, recombination or region-specific rates of evolution (Gorbalenya & Lauber, 2022). Moreover, VirusTaxo was only tested to predict novel (unseen) genomes by removing one species from the training dataset and predicting it at the correct genus. However, the sequence similarity is usually very high among members of the same genus (more than 96% in the tested case).

Different approaches using CNN for DNA sequences in various fields achieved high accuracy, even in the identification of viruses, as seen in DeepVirFinder, but there is a lack of viral taxonomic classification tools using 2D-CNNs. This reinforces the need for the development of additional tools for viral taxonomic classification.

One interesting group of viruses that were shown to have greater diversity than previously thought are the DNA viruses with small, circular, single-stranded genomes. These viruses do not seem to cause disease in humans but are known to cause major losses in crops and livestock, and in consequence, have been studied in detail through the years (Rosario *et al*., 2012). Among these viruses, some families comprise a distinct group that encode a conserved replication-initiation protein (named Rep) involved in the initiation step of rolling-circle replication, an important replication mechanism for circular DNA genomes also used in some plasmids. These families were informally named CRESS-DNA (<u>c</u>ircular, <u>Re</u>p-encoding <u>ssDNA</u>) viruses, and were recently classified in a new phylum designated *Cressdnaviricota* (Krupovic *et al*., 2020). The family *Geminiviridae*, which includes a large number of economically important plant viruses that cause severe diseases in tropical and subtropical reagions worldwide (Rojas *et al*., 2018), is classifed in this phylum.

The objective of this work was to evaluate the capacity of 2D-CNNs to be used in the taxonomic classification of viruses, using members of the phylum *Cressdnaviricota* as test subjects.

## MATERIALS AND METHODS

### Computational resources

All code and analyses in this work were done using Python 3 (python.org/doc/). All models and functions used are from Keras (github.com/keras-team/keras), TensorFlow (TF, (Abadi *et al*., 2016) and SKlearn (Pedregosa *et al*., 2011).

### Data collection

All sequences used in the training of the model were obtained from the ICTV’s Virus Metadata Resource (VMR, ictv.global/vmr), a taxonomic list of well-characterized virus isolates from all classified species, using the 08/31/2022 release (VMR_20-190822_MSL37.2.xlsx). From this file, GenBank accession numbers of all *Cressdnaviricota* members were extracted and used to download the genomic sequences from the database using Entrez, available in the Biopython library (Cock *et al*., 2009). Also, as the final test dataset for benchmarking the model against VirusTaxo, all *Cressdnaviricota*-associated information available was downloaded from the NCBI genome data hub (ncbi.nlm.nih.gov/data-hub/genome/) using Entrez, and those accessions that were in the training set or reference sequences from NCBI (started with “NC_”) and had 100% Blast identity and coverage with model sequences were removed.

### Convolutional Neural Networks

Based on the idea of visual perception, CNNs were created from artificial neural networks (ANN) to extract features from data. CNNs have been used mainly for image recognition using 2D convolutional layers, but also for text, using 1D convolutional layers (Li *et al*., 2022). In our classification problem, we will be testing supervised (labeled data) 2D-CNNs, which are commonly composed of different types of layers and where each one may change the data in specific ways. Most of the times they are composed of one input layer that receives matrices of fixed dimensions, one or more hidden layers, and one output layer with the prediction. In our classification problem, the output layer produces a fixed-size vector that has the same size as the number of categories/taxa being classified in the model, where each position in the output vector defines (using the softmax activation function for classification, described by Araújo *et al*. (2017) the probability of the tested sample to belong to each class, and where the sum of all positional values is equal to 1.

Training occurs by updating the network trainable parameters by applying a stochastic gradient and back-propagation to reduce loss (LeCun *et al*., 2015), which in our case was the Adam optimizer (Kingma & Ba, 2014) with default parameters and the categorical cross-entropy loss function, which “evaluates the difference between the probability distribution obtained from the current training and the actual distribution” (Li *et al*., 2022), and that is often applied in the case of multiclass classification. We tested different architectures with common layers available to use in 2D-CNNs with TF: convolutional, pooling, dropout and dense (also called fully connected) layers, in a “manual search”, trying each architecture with hyperparameters similar to those found on CNN sequence classifiers that yielded good results, and then refining them by testing variations and combinations.

### Convolutional layers

Convolutional layers are the main layers in 2D-CNNs. Each convolutional layer is made of a user-selected number of filters, commonly called kernels, which are matrices with specified sizes (e.g. a 2D kernel/filter as a 3x3 matrix; Figure 1). As happens in neurons of dense layers, each kernel from a convolutional layer is normally initialized with random numbers in each of its values in the matrix (called weights, which are trainable parameters) and will move through the input matrix (like a sliding window), calculating the dot product, adding bias (which also are trainable parameters), and applying a nonlinear activation function (which avoids direct linearity between input and output), which in our test was the rectified linear unit (ReLU) (Nair & Hinton, 2010), where numbers lower than zero are converted to zero, and the rest is directly used as output. In the end, each filter produces a result matrix (also called features map) as output for the next layer (Li *et al*., 2022).

**Figure 1.**
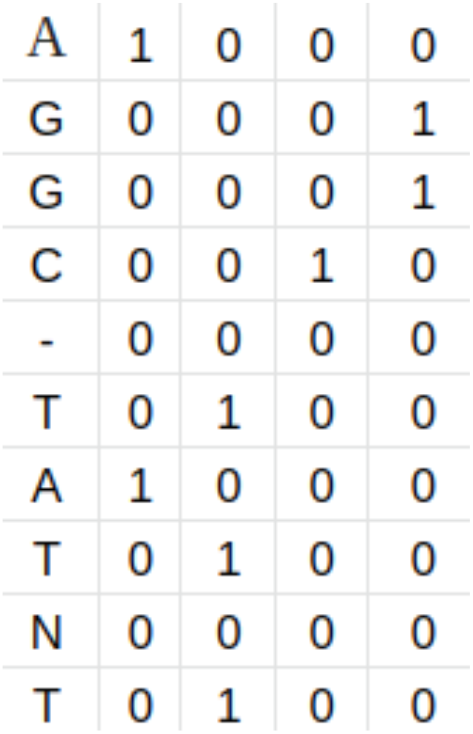
Example of one-hot encoding for the sequence AGGC-TATNT, where A = [1,0,0,0], T = [0,1,0,0], C = [0,0,1,0], G = [0,0,0,1] and any other character = [0,0,0,0].

Besides a number of filters and their size, other important parameters in the convolutional layer are padding and stride. Padding, when selected to “same” as used in this work, consists in adding zeros (or other values) around the sides of the image/matrix, allowing the convolution kernel to take the borders’ values into consideration more than once and thus reducing the boundary effect (some filter sizes and strides could lead to edges being less used, which has been shown to affect the output) (Islam *et al*., 2021). Stride is defined as the number of steps that the kernel will move (“slide”) in the input matrix for each convolution (we used the standard, where the stride is equal to 1). One example can be seen in Figure 1, where the 3x3 kernel applies convoluting in the first three rows and first three columns of the matrix. After that, the kernel will move two columns (stride defined as 2 or 2x2) to the right and repeat the convolution, repeating this action until the end of the padded matrix, and then going back to the left side of the input matrix, but two lines down (again stride 2), starting again and repeating until the end of the input matrix (Li *et al*., 2022).

### Pooling layers

The pooling layer reduces the dimensions of an output matrix from the convolutional layer, thus reducing the need for computational resources. Different strategies can be used, such as average-pooling or max-pooling, by merging features. When using max-pooling, only the highest value inside the pooling window is propagated to the next layers, e.g. a max pooling layer with a 2x2 window size will reduce the input matrix dimensions by a factor of 2, so that a 4x4 matrix will be reduced to a 2x2 matrix (LeCun *et al*., 2015).

### Dropout layers

These are regularization techniques that help reduce overfitting (when the model “memorizes” the training dataset instead of finding a predictive rule, not working well in unseen data) (Dietterich, 1995), in this case by dropping a defined percentage of random connections/input values, e.g. a 0.3 dropout will cause 30% of the connections to be ignored (Srivastava *et al*., 2014).

### Flatten and dense layers

The output matrix from a 2D convolution and max pooling layers may be then passed to the flatten layer, which will turn all output matrices received into a single concatenated vector to be used as input at the dense layers. Each dense layer is composed of a selected number of artificial neurons, where each neuron receives as input all features/neurons outputs from the previous layer, then performs multiplication of each input value by a specifically position-related weight, sums all multiplications and adds the neuron “bias” (also known as neuron internal state). This value goes through the activation function (ReLU again in our model), which simulates the activation of a biological neuron by modulating the information to be released as output to the next layer. In the last layer, in our classification case, the number of categories in the model is used as the number of neurons, where a softmax activation function is used to get a vector of probabilities for the current sample to belong to each category. While training, the difference between the predicted taxonomic group and the true group is computed by the loss function and back-propagated through the network, updating the weights and biases of the neurons in each layer (Li *et al*., 2022).

### Early stopping

Also for dealing with overfitting, early stopping is based on detecting when overfitting starts during training by monitoring a metric such as accuracy or, in our case, validation loss, to stop training when the loss reaches a minimum for more than a fixed number of epochs (the patience), and recover the best epoch (smaller loss) model parameters. This technique has been widely used for its simplicity and ease of implementation (Prechelt, 2012). We applied it as described in the TensorFlow documentation (github.com/tensorflow/docs).

### Evaluation metrics

For performance evaluation, the most commonly used metric for balanced datasets is arguably the Area Under the Receiver Operating Characteristic (ROC) Curve (AUC-ROC) (Melo, 2013). When dealing with imbalanced datasets, the Area Under the Precision-Recall Curve (AUC-PR) is a better metric, as it takes into account the precision parameter (Sofaer *et al*., 2019). These metrics are based on the number of true positives, true negatives, false positives and false negatives. From them, the true positive rate (y-axis) and false positive rate (x-axis) are calculated for the ROC curve, and the recall (x-axis) and precision (y-axis) are calculated for the PR curve (Davis & Goadrich, 2006). Another useful metric for classification is the F1 score, which is the harmonic mean of precision and recall (Wardhani *et al*., 2019). The F1 score is more affected by the classification itself then the prediction probabilities. In our case, for multiclass classification during training, which frequently deals with imbalance in the datasets, we applied the AUC-PR metric available on Keras (keras.io/api/metrics/classification_metrics). After training, the F1 score (weighted for each group using sklearn libraries) and the AUC for the model were obtained for the current validation dataset.

### Cross-validation

When evaluating the performance of models in ML, the common use of a training set and a test set also avoids overfitting, by training the model in part of the dataset and evaluating its performance in unseen data, guaranteeing that the model did not memorize each label. This practice is known as cross-validation (CV) and is well established for evaluating the model’s response to different hyperparameters. One commonly used method of CV is known as K-Fold, where the dataset is split into *K* smaller sets of data, and each subset is used as the test set one time, while the remaining data is used as the training set. The process is repeated *K* times with each subset as the test set, and the metrics are used to obtain an average of the model’s performance (Hastie *et al*., 2013). We applied this method using the available Keras function with *K* = 5 during hyperparameter tuning and also using its stratified version, which splits the data based on a percentage of each class, for dealing with imbalanced datasets.

### Data encoding and input standardization

One-hot encoding is a simple and practical way of encoding textual data in a numerical format, used in different computation approaches. In DNA and protein codification, each possible base/amino acid is converted to a fixed size vector, with zeros in all positions except in one (hence the method’s name), corresponding to that letter (base or aminoacid letter symbol) in that position. In our case, for DNA we codified the nucleotides A, T, C and G as [1,0,0,0], [0,1,0,0], [0,0,1,0] and [0,0,0,1], respectively, while for every other possible DNA representation we used the vector [0,0,0,0]. So, for example, when encoding a DNA sequence of length L, the algorithm transforms the sequence in a Lx4 matrix, where each line represents the position of each DNA base in the sequence and each column represents which letter is using “ones” (Figure 1).

In most works this matrix is applied to a 1D-CNN, where each nucleotide position (column) goes through individual channels which search for patterns within each type of nucleotide. In our approach, we use the less common option of using this matrix as an image and passing it through a single-channel 2D-CNN. Thus, the matrix is used as one whole thing, with each filter seeking patterns in all four nucleotides at the same time.

Before the coding part of the sequences, an important step is formatting the input sequences. Initially, as some viruses have multipartite genomes (where their genomic information exists in multiple segments encapsulated in different viral particles), we concatenated the segments into a single molecule, as CNNs kernels are not positionally limited in consequence of moving entirely through the matrices looking wherever the patterns may be. Moreover, CNNs require a fixed-size matrix as input, which has been done differently by each approach, like completing with zeros/gaps the remaining size or truncating the sequences in specific sizes. In our case, seeking to use complete genomes in all inputs, this was done based on the idea seen in copy-paste augmentation methodology for image recognition, where copies of one object are added in the same image, and which has been shown to be a very effective and robust augmentation method (Ghiasi *et al*., 2021; Pappagari *et al*., 2021). Thus, every sequence had its size completed to the fixed input length by adding copies of itself to its end portion until it reached the desired length, which in this work was set to 15,000 positions/nucleotides (which means using a 15,000 x 4 matrix as input). In the case of the longest *Cressdnaviricota* genomes, those of viruses from the genus *Nanovirus* (8 segments with aproximately 1,000 nucleotides each), this lead to almost two copies of the concatenated genome.

### Data imbalance

The composition of the dataset is arguably the most important part when developing a prediction model, since the model’s ability to predict is defined and limited by the dataset. The dataset is considered to be imbalanced when the different categories have data distributions that are considerably different amongst themselves. For example, if almost all samples in a dataset with two groups belong to one of them, the model could get biased in classifying every sample as the largest group, as it would get good accuracy by that. To solve this problem, different methods have been proposed, mostly based on different ways of undersampling the largest dataset or oversampling the smallest dataset (Lin *et al*., 2017; Masko & Hensman, 2015).

In our study case there is an imbalance in different taxonomic ranks, mainly caused by the bias on classifying economically important pathogens. The main example is the genus *Begomovirus,* belonging to the family *Geminiviridae*, which includes several hundred species, with more than 600 genomes in the VMR file. Seeking to reduce this number, we applied CD-Hit (Fu *et al*., 2012) to cluster all sequences with more than 70% sequence identity, with other parameters set to default. After this clustering, we only used one sequence as representative of each cluster, reducing the number of sequences from more than 600 to 118.

We also checked the effect of two techniques to solve oversampling during the tests, trying to balance small datasets (such as at the genus level, where many genera only have a small number of sequences associated with them) by creating variations in copies of the existing data. In the first tecnique, mutations of a random percentage of bases are applied to a specific number of copies of existing sequences, similar to what is suggested by Busia *et. al.* (2020), randomly mutating 1-2% of different sequences until the groups are equivalent in data. The second approach was the addition of a sequence fragment, randomly selected from a pool, to each border of sequence copies, e.g. inittialy splitting the sequences of each group into fragments that are held in the same pool and are randomly sampled. This is also similar to the idea of copy-paste augmentation, but with fragments of the viral sequences being added to the borders to increase the data variability. Considering that most viruses have compact genomes and most portions of the genome normally carry information associated with that sequence, it is unlikely that the fragments will not carry new information/patterns to the sequence.

### Hyperparameter tuning

After defining the main methodologies, functions and metrics, important hyperparameters (number and types of layers, number of kernels, kernel size, dropout, max-pooling windows size, number of neurons in dense layers) were evaluated by testing some previously used architectures for DNA sequence classification with CNNs. In one recent example (Dasari & Bhukya, 2022), a CNN model for the detection of human viral pathogens on human metagenomic datasets applied a 1D-CNN architecture composed of three groups of layers for extracting more complex features in every layer. Each group of layers applied sequentially: a 1D convolutional layer (32 kernels, kernel size 7; 8 kernels, size 4; 8 kernels, size 3), a dropout layer (0.2 in all three groups), and a max-pooling layer (kernel size 2, stride 2 in all three groups). Then, the flattening layer condensed the information into a vector used as input to a dense layer with 32 neurons. Finally, an output layer with two neurons released the classification (viral or non-viral sequence) with 99% accuracy on training.

A different approach was applied with good results by Min *et al*. (2021) using larger window sizes. In this case, the 1D convolutional layer and the max-pooling layer used filter sizes ranging from 40 to 60 and from 20 to 30, respectively. These larger kernels are interesting because they appear to be similar to the optimal size of k-mers used in VirusTaxo, which is 21 nucleotides. Also when converted into nucleotide sequences, the sizes of the average canonical protein backbone fragments used in databases (4 to 14 amino acids) (Baeten *et al*., 2008) would correspond to a range of 12 to 42 nucleotides. Based on these different architectures, we tried applying them directly, combined, and with different variations as seen in other studies.

## RESULTS AND DISCUSSION

### Model architecture

For testing the different possible model architectures, we initially used as datasets the first branching of *Cressdnaviricota*, that between the classes *Alfiviricetes* and *Repensiviricetes.* This allowed us to test without using augmentation techniques, as the proportion of members between the groups is almost 1:2 (252 and 474 members for *Alfiviricetes* and *Repensivirices*, respectively), which other CNN studies have shown not to be detrimental to the results (Qu *et al*., 2020). As this dataset comprises all sequences that would be used for training and are defined in only two groups, it seemed practical for evaluating the performance effect of different architectures.

The performance was evaluated during training using AUC-PR applying the *K*-fold approach (*K* = 5), where for each fold the AUC-PR and the F1 score were calculated based on the validation set and used to get the final average. Also, the early stopping function was set to monitor the validation loss, and the patience was set initially to 5. However, after testing higher values, we found that even after some dozens of epochs the loss could still drop and classificate more sequences correctly. So from almost the beginning we set the patience to 100, with a higher number of maximum epochs to accommodate large trainings, and started getting better results after a few hundred epochs. Finally, we initially used a batch size of 100, but in the first trials we found that using a lower value tended to improve the accuracy so it was eventually set to 2, which gave better results and agrees with the results of Masters & Luschi (2018).

The first architecture tested was the one used by Dasari & Bhukya (2022). We applied it directly, adapting only the filter sizes to 2D convolution (7x7, 4x4 and 3x3 convolutional kernel sizes, and 2x2 max-pooling sizes). This configuration achieved an average AUC-PR of 93.131 with standard deviation (STD) of 1.498, and average F1 score of 86.662 and STD of 3.302, which meant fewer than a hundred sequences being wrongly classified. Nevertheless, the validation AUC and especially the loss graphs are highly scattered (Figure 2), indicating that this architecture cannot find distinguishing patterns in the data.

**Figure 2.**
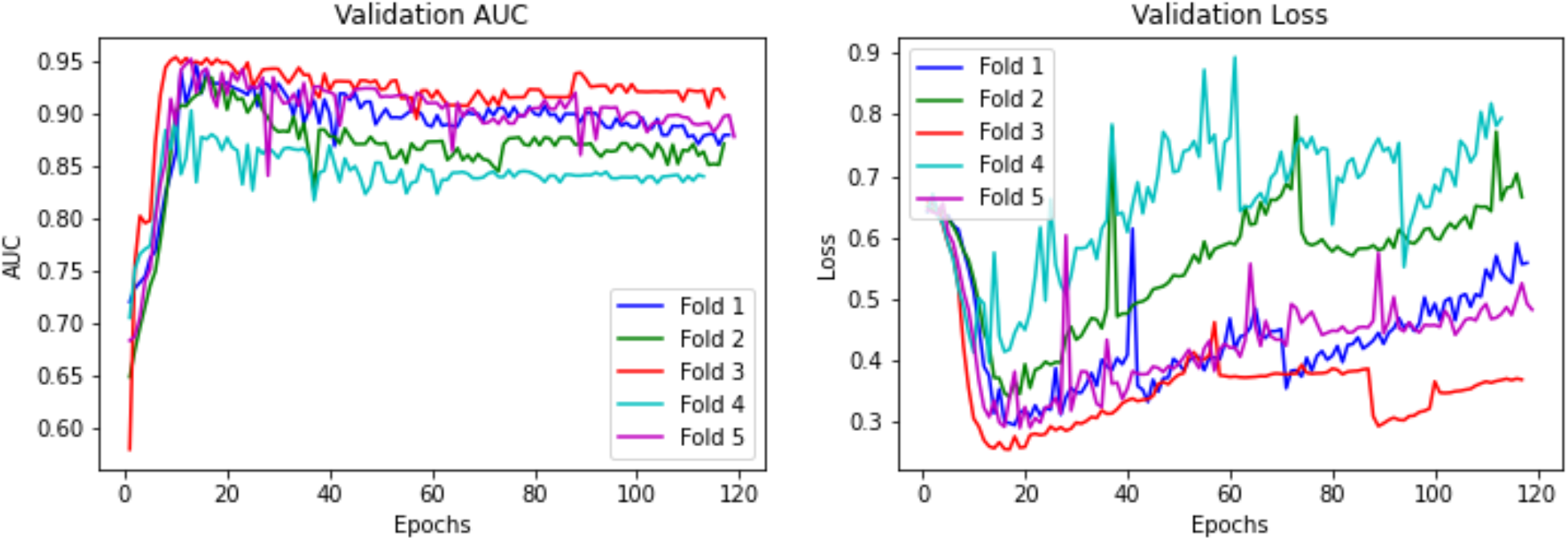
Training metrics (validation AUC-PR and loss) of a 2D-CNN for sequence-based viral classification in the phylum *Cressdnaviricota*, using a similar architecture than Dasari & Bhukya (2022).

Besides other changes that did not affect the results, we tried using the larger filter sizes for convolutional and max-pooling layers, as proposed by Min *et al*. (2021). Thus we changed the sizer of all three layers using 40x4 filters (only four columns as the sequence vector has only four positions) in the first two convolutional filters, and 30x4 in the last one, and changing the max-pooling filter to 20x4, seeking to find higher patterns in convolutions and reduce the dimensions, keeping only what is important (higher values). With these changes the model achieved an average AUC-PR of 99.4 (STD of 0.547) and average F1 score of 96.774 (STD of 2.585) (Figure 3), with only two dozen sequences being wrongly classified, thus showing that larger filters increased the prediction power of the CNN. However, even with good AUC and F1 scores the model made wrong predictions, indicating that there was still space for further improvement.

**Figure 3.**
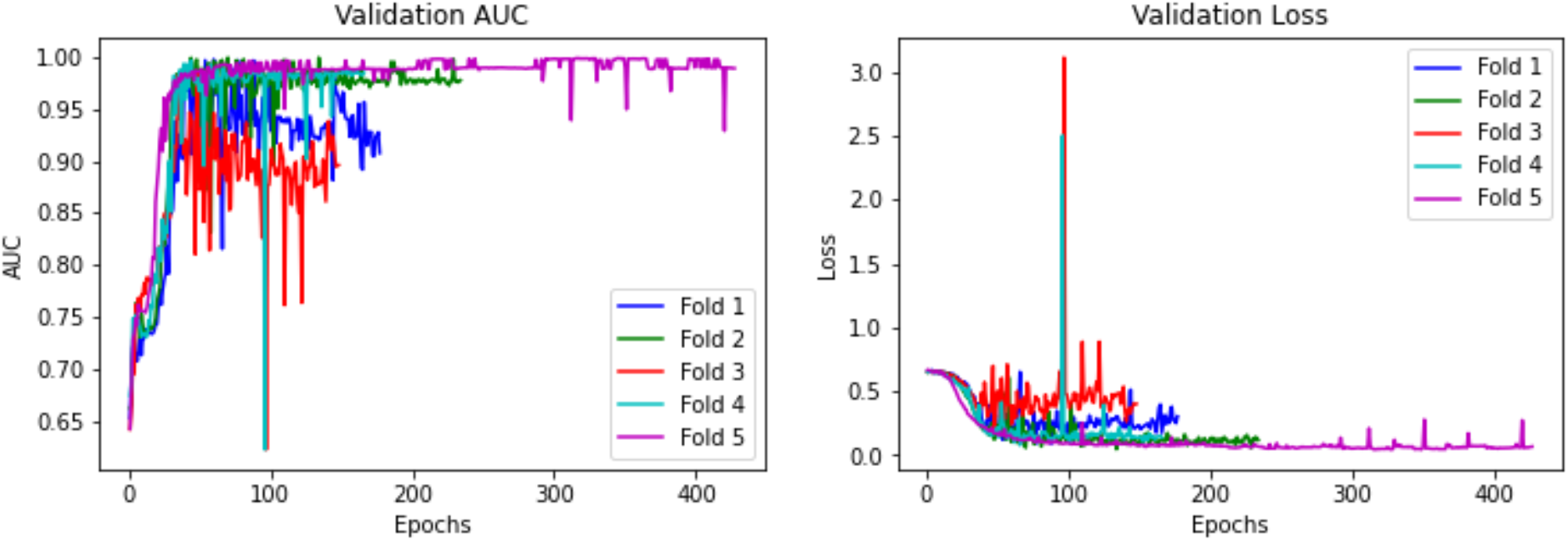
Training metrics (validation AUC-PR and loss) of a 2D-CNN for sequence-based viral classification in the phylum *Cressdnaviricota*, using the architecture proposed by Dasari & Bhukya (2022) with larger filters (convolutional kernels, 40x4 and 30x4; max-pooling, 20x4) as proposed by Min *et al*. (2021).

We then tried different combinations of hyperparameters, changing the number of layers, the number of convolutional kernels and their sizes, layer orders, dropout percentages, and max-pooling filter sizes, changing them one at a time, maintaining those configurations that enhanced the model’s performance and trying additional variations from them. After extensive testing with different architectures, an excellent performance was achieved by reducing the max-pooling column size from 4 to 1 (where the dimensions are merged for each nucleotide, maintaining the matrix width). This configuration achieved an average F1-score of 98.893 (STD of 0.709) and AUC-PR of 99.762 (STD of 0.364). The validation graphs (Figure 4) show that the curves quickly attained good values and started flattening, with more punctual peaks compared to the previous configurations, indicating more stability during training.

**Figure 4.**
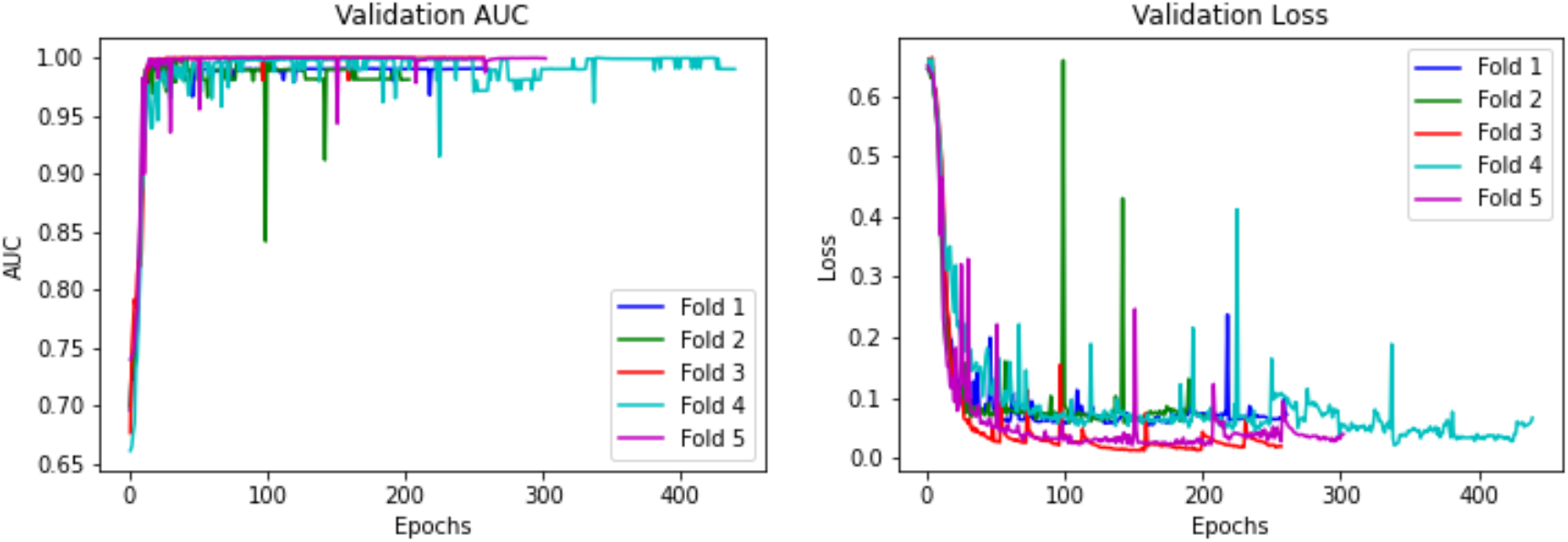
Training metrics (validation AUC-PR and loss) of a 2D-CNN for sequence-based viral classification in the phylum *Cressdnaviricota*, using an architecture incorporating elements from Dasari & Bhukya (2022) and Min *et al*. (2021) and the width of max-pooling layers reduced from 4 to 1.

Another change that enhanced the model’s performance was removing one group of convolutional layers and changing the convolutional kernel size. The first CNN that stood out used 16 150x4 filters in the first layer, followed by a max-pooling layer with a 50x1 filter and a dropout layer of 0.5. The next convolutional layer had 32 50x4 filters, max-pooling of 25x1 and dropout also of 0.5. This was followed by the flatten layer, a dense layer with 64 neurons, another dropout layer with 0.4, and finally the output layer releasing the prediction vector. With this architecture the model was able to achieve an average AUC-PR of 99.812 (STD of 0.369) and F1 score of 99.862 (SDT of 0.275), with only one sequence wrongly predicted. The validation graphs (Figure 5) display patterns of flattening and punctual peaks, however more uniform between the five folds, and also with values of AUC varying mostly near 0.999. The loss graph was more constant between folds, with considerable overlapping.

**Figure 5.**
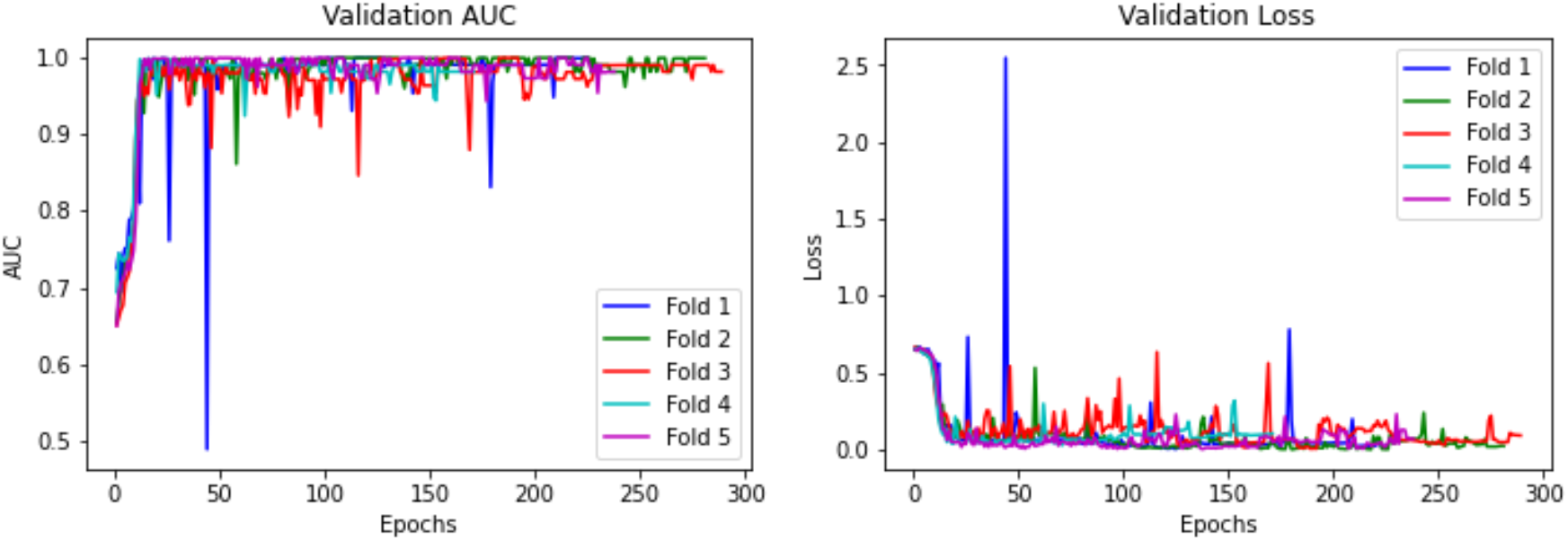
Training metrics (validation AUC-PR and loss) of a 2D-CNN for sequence-based viral classification in the phylum *Cressdnaviricota*, using an architecture incorporating elements from Dasari & Bhukya (2022) and Min *et al*. (2021), but with the width of max-pooling layers reduced from 4 to 1, one less group of layers, and bigger filter sizes.

A final change in the model (64 instead of 16 convolutional kernels in the first convolutional layer; Figure 6) achieved perfect (1.0) AUC and F1 scores, with 0 STD and no wrongly classificated sequences. A slightly worse result (two wrongly classified sequences) was obtained in a sequential test.

**Figure 6.**
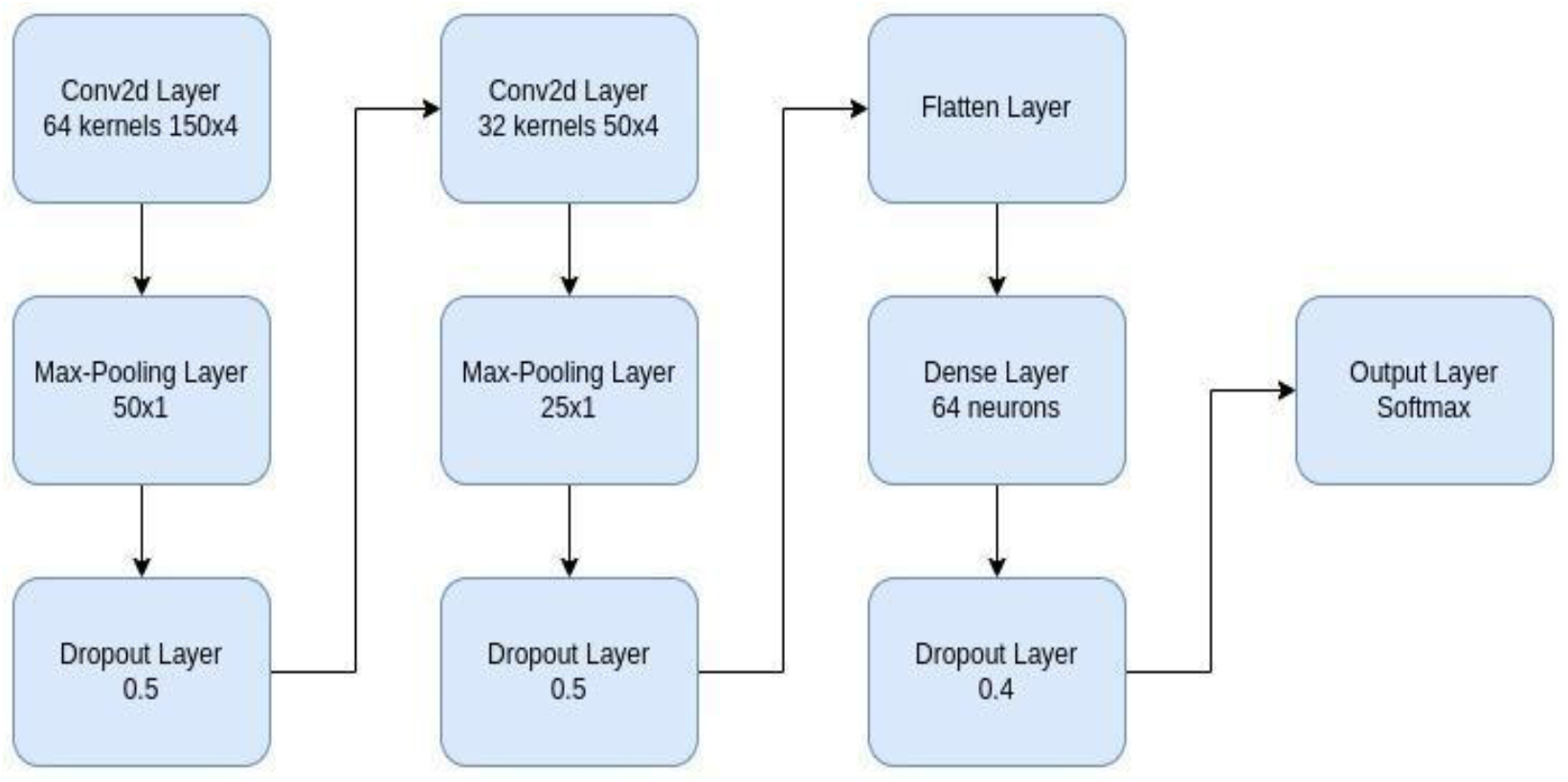
Architecture of the CNN with best results during training, where in one test it classified all sequences correctly.

With these high value metrics for the two latter architectures, we chose to use the one with 64 initial convolutional kernels to test for lower taxonomic ranks. First we trained the model for the family *Circoviridae*, which includes two genera, *Circovirus* and *Cyclovirus*, with 49 and 52 sequences, respectively. The model achieved high values, with an average AUC-PR of 99.902 (STD of 0.120), and F1 score of 97.995 (STD of 2.456), indicating that it worked well in balanced data even for the lower ranks.

However, imbalanced data composes most of the families in the phylum, so we also trained the model in one imbalanced family of metagenomic characterized viruses, the *Genomoviridae*. In this case the model achieved an AUC-PR of only 84.209, while the F1 score was 89.369. The model made mostly correct predictions but with low confidence in the probabilities, with 24 wrongly classified sequences in a dataset of almost 100 sequences. The low AUC-PR and F1 scores can be expected considering that one genus has only three sequences, seven genera have about 10 sequences in average, one genus has 50+ sequences, and one genus (*Genomovirus*) has 190+ sequences. This imbalance, which is also observed in most of the other families and is a common sampling problem in virus taxonomy, is likely the main factor responsible for the lower values obtained for the two metrics.

One important characteristic of ML algorithms is the division of the dataset into at least two groups, one for training and one for testing. Thus, when only one sequence is available in a genus (known as singletons), this genus was removed from the training set. This was the case for 26 genera. To avoid ignoring these genera, each input was also analyzed by BLAST (Altschul *et al*., 1990) against these singletons.

### Pipeline development

We decided to use a hierarchical pipeline for classification which is based on a series of models parsing each input by classifying them from the uppermost rank of phylum down to genus, which means predicting four taxonomic ranks (or three in some cases, see below). This hierarchical methodology seeks to reduce the sampling problem shown by the small number of species in some genera, trying to confine the bias to the related family model with imbalance.

The pipeline first receives an input matrix from each sequence by one-hot encoding (as described in the section *Data encoding and input standardization* in the Material and Methods). This matrix is then used for prediction at all ranks, where the predicted rank is used to select the model to be used for the next rank prediction. For example, the first model separates the *Cressdnaviricota* members between those classified in the classes *Arfiviricetes* or *Repenviricetes,* and marks the prediction type as done by “model”. Once the class has been predicted, the matrix is then used to obtain the order predictions by the second model (in case of only one existing group there is no model, thus the prediction type is marked as “single” and goes directly to the existing rank), repeating this process until reaching the genus classification. After the genus prediction, the pipeline runs a BLAST analisys against all sequences from the other genera in the predicted family, checking if only sequences from the predicted genus have alignment similarity. When there are BLAST hits for the other genera (including the singletons), it uses the hit with the highest identity and coverage as the output prediction.

### Pipeline benchmarking

The taxonomic classification capability of our model was first evaluated by comparing with that of VirusTaxo to classify *Cressdnaviricota* sequences available on the NCBI’s genome datahub. After removing the used accession numbers and reference sequences that had 100% identity and coverage using BLAST, we selected only the sequences that were classified at the genus level, which left us with 4,170 sequences for testing. For comparison with VirusTaxo, we used its database creation function to build a custom dataset with all sequences used in our model training, using a k-mer size of 21, which is the value indicated by the program for DNA sequences.

The models tested here were built using the best architectures found above with two augmentation techniques (percentage augmentation, PercAug, and border augmentation, BordAug), with no augmentation (NoAug), and also with a combination of the phylum model using NoAug while the other models used BordAug or PercAug, totaling 10 different pipeline ensembles tested against VirusTaxo. Also, after the predictions, all results were parsed and sequences that had 100% identity and coverage were again removed, which lead to 4,051 sequences remaining that were never “seen” by the models. The vast majority (3,647) of these sequences belong to the genus *Circovirus*, while some other genera had only one or two sequences, such as *Curtovirus*, *Grablovirus*, *Capulavirus*, *Topilevirus*, *Bacilladnavirus*, *Gemykolovirus* and *Gemykrogvirus*. As a consequence of this small number of sequences, these genera and some others with few members (or that were misclassified by all tests) were not placed in the final result calculations below.

All tested pipelines got metrics that were similar to those obtained with VirusTaxo for the most populated genera, but most of them misclassifed fewer sequences then VirusTaxo (Table 1, Table 2). The pipeline composed of a NoAug model for phylum and BordAug for the remaing models achieved the highest accuracy (98.56%) and only misclassified 58 viruses.

**Table 1.**
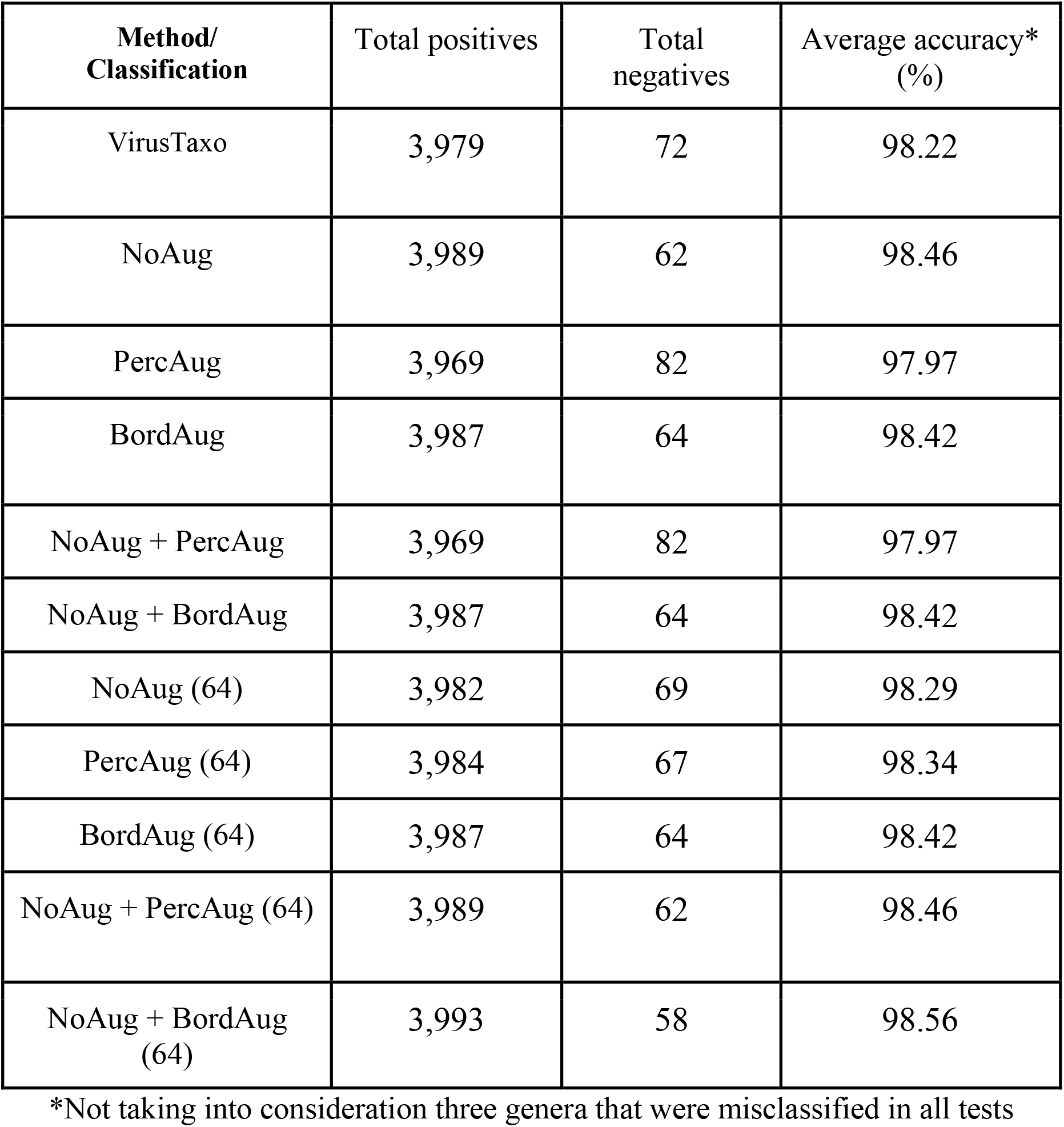
Classification results for the benchmarking dataset tested with 10 different model ensembles developed here and with VirusTaxo, where positives are sequences classified correctly, while negatives represented the misclassified sequences.

| **Method/**<br>**Classification** | Total positives | Total<br>negatives | Average accuracy*<br>(%) |
| --- | --- | --- | --- |
| VirusTaxo | 3,979 | 72 | 98.22 |
| NoAug | 3,989 | 62 | 98.46 |
| PercAug | 3,969 | 82 | 97.97 |
| BordAug | 3,987 | 64 | 98.42 |
| NoAug + PercAug | 3,969 | 82 | 97.97 |
| NoAug + BordAug | 3,987 | 64 | 98.42 |
| NoAug (64) | 3,982 | 69 | 98.29 |
| PercAug (64) | 3,984 | 67 | 98.34 |
| BordAug (64) | 3,987 | 64 | 98.42 |
| NoAug + PercAug (64) | 3,989 | 62 | 98.46 |
| NoAug + BordAug<br>(64) | 3,993 | 58 | 98.56 |

\*Not taking into consideration three genera that were misclassified in all tests

**Table 2.** Accuracy results for each genus using the benchmarking dataset, tested with 10 different model ensembles developed here and with VirusTaxo, currently the tool with higher accuracy for virus taxonomy.

| **Method/**<br>**Classification (%)** | *Circovirus*<br>(3,647) | *Begomovirus*<br>(118) | *Gemycircularvirus*<br>(42) | *Cyclovirus*<br>(17) | *Porprismacovirus*<br>(31) | *Nanovirus*<br>(10) | *Babuvirus*<br>(125) | *Mastrevirus*<br>(8) | *Gemykibivirus*<br>(18) | *Huchismacovirus*<br>(23) |
| --- | --- | --- | --- | --- | --- | --- | --- | --- | --- | --- |
| VirusTaxo | 98.43 | 100.0 | 80.95 | 94.11 | 93.54 | 90.0 | 100.0 | 87.5 | 94.44 | 100.0 |
| NoAug | 98.71 | 100.0 | 80.95 | 94.11 | 90.32 | 90.0 | 100.0 | 100.0 | 94.44 | 95.65 |
| PercAug | 98.16 | 100.0 | 80.95 | 88.23 | 93.54 | 90.0 | 100.0 | 100.0 | 94.44 | 95.65 |
| BordAug | 98.62 | 100.0 | 80.95 | 88.23 | 96.77 | 90.0 | 100.0 | 100.0 | 94.44 | 95.65 |
| NoAug<br>+PercAug | 98.16 | 100.0 | 80.95 | 88.23 | 93.54 | 90.0 | 100.0 | 100.0 | 94.44 | 95.65 |
| NoAug +<br>BordAug | 98.62 | 100.0 | 80.95 | 88.23 | 96.77 | 90.0 | 100.0 | 100.0 | 94.44 | 95.65 |
| NoAug (64) | 98.46 | 100.0 | 80.95 | 100.0 | 93.54 | 90.0 | 100.0 | 100.0 | 94.44 | 95.65 |
| PercAug (64) | 98.54 | 100.0 | 80.95 | 94.11 | 93.54 | 90.0 | 100.0 | 100.0 | 94.44 | 95.65 |
| BordAug (64) | 98.60 | 100.0 | 80.95 | 100.0 | 93.54 | 90.0 | 100.0 | 100.0 | 94.44 | 95.65 |
| NoAug +<br>PercAug (64) | 98.68 | 100.0 | 80.95 | 94.11 | 93.54 | 90.0 | 100.0 | 100.0 | 94.44 | 95.65 |
| NoAug +<br>BordAug (64) | 98.76 | 100.0 | 80.95 | 100.0 | 93.54 | 90.0 | 100.0 | 100.0 | 94.44 | 95.65 |

The results also demonstrate how augmentation by itself had a small and not guaranteed beneficial effect on the results, with a worse result in one case, and how by mixing methodologies it was possible to achieve better metrics. Another noticeable result is that some sequences previously classified as circoviruses in the NCBI database got higher identity and coverage with sequences in the genus *Cyclovirus*. These sequences were classified as mistakes, but with the possibility that the error actually took place in the original classification, as they did not get BLAST hits with the circoviruses used in the model. Furthermore, all sequences in the genera *Drosmacovirus* and *Glamdringvirus*, and eight sequences in the genus *Gemycircularvirus* were always wrongly classified, which suggests that these samples may have very different sequences from the ones used in the training dataset or need to be reclassified.

## CONCLUSIONS

This work has shown again that CNNs have a really good capacity to differentiate between samples and are able to find conserved patterns in genomic sequences, allowing their use in taxonomic classification. We tested two methodologies for augmentation, where adding fragments of sequences to the borders of each new sequence often yielded better results than with no augmentation. Conversely, adding a percentage of mutation to the sequence yielded worse results in some cases, which could be consequence of divergent/deleterious mutations made in regions of important patterns during augmentation.

Furthermore, it was possible to confirm that the randomness that exists on the hyperparameters tunning changed the results from one trained model to the other, even maintaining the same architecture, suggesting that creating and comparing models, even using different augmentation techniques, could lead to different, and often better, results. Here we present a novel method for classifying viruses in the phylum *Cressdnaviricota* down to the genus level using a convolutional neural network (CNN), and achieving highly accurate results.

